# Consistent specificity and efficiency of tamoxifen-mediated cre induction across ages

**DOI:** 10.1101/2023.09.19.558482

**Authors:** Collyn M. Kellogg, Kevin Pham, Sunghwan Ko, Jillian E. J. Cox, Adeline H. Machalinski, Michael B. Stout, Amanda L. Sharpe, Michael J. Beckstead, Ana J. Chucair-Elliott, Sarah R. Ocañas, Willard M. Freeman

**Affiliations:** Genes & Human Disease Program, Oklahoma Medical Research Foundation, Oklahoma City, OK USA; Department of Biochemistry & Molecular Biology, University of Oklahoma Health Sciences Center, Oklahoma City, OK USA; Aging & Metabolism Program, Oklahoma Medical Research Foundation, Oklahoma City, OK USA; Department of Pharmaceutical Sciences, University of Oklahoma Health Sciences Center, Oklahoma City, OK, USA; Oklahoma City Veterans Affairs Medical Center, Oklahoma City, OK USA; Neuroscience Graduate Program, University of Oklahoma Health Sciences Center, Oklahoma City, OK USA; Department of Physiology and Neuroscience, University of Oklahoma Health Sciences Center, Oklahoma City, OK USA

## Abstract

Temporally controlling cre recombination through tamoxifen (Tam) induction has many advantages for biomedical research. Most studies report Tam induction at early post-natal/juvenile (<2 m.o.) mouse ages, but age-related neurodegeneration and aging studies can require cre induction in older mice (>12 m.o.). While anecdotally reported as problematic, there are no published comparisons of Tam mediated cre induction at early and late ages. Here, microglial-specific Cx3cr1^creERT^^2^ mice were crossed to a floxed NuTRAP reporter to compare cre induction at early (3-6 m.o.) and late (20 m.o.) ages. Specificity and efficiency of microglial labeling at 21-22 m.o. were identical in mice induced with Tam at 3-6 m.o. or 20 m.o. of age. Age-related microglial translatomic changes were also similar regardless of Tam induction age. Each cre and flox mouse line should be validated independently, however, these findings demonstrate that Tam-mediated cre induction can be performed even into older mouse ages.

## Introduction

The development of the cre-LoxP system^1,2^ with subsequent integration^3^ and refinement^4^ of a cre bound to a mutated estrogen receptor (ERT2) allows for temporally controlled induction of cre recombination with tamoxifen (Tam) administration^4–6^. When cre-ERT2 expressing mice are crossed to mice with LoxP sites flanking a stop codon or desired gene, induction of the cre recombinase via Tam injection activates or suppresses the transgene, respectively^7,8^. Including a gene promoter of a cell type-specific gene upstream of the cre-ERT2 gene can combine cell specificity and temporal control^9,10^. This system has been widely adopted and utilized across biomedical research. While a plethora of reports describe the validation of different cre- ERT2 expressing mouse lines and examine the effects of altered transgene expression, most studies report induction of cre recombination at young/adolescent age (<2-3 m.o.).

In aging research and age-related neurodegeneration studies, experimental designs that induce cre recombinase at old ages can be advantageous. A longstanding hypothesis in aging research is that expression of certain genes is positively adaptive at young age until late age, where it becomes maladaptive^11^. In neurodegeneration studies, neuroinflammatory responses may be prodromally beneficial to contain disease spread but deleterious over longer periods, contributing to disease progression^12–17^. Thus, experimental designs where Tam induction of cre recombinase is performed in older mice are warranted. However, to our knowledge, there are no reports comparing *in vivo* cre induction in young and old mice.

Tam is metabolized by cytochrome P450 into the metabolites N-desmethyltamoxifen, 4- hydroxytamoxifen, endoxifen, and norendoxifen, with 4-hydroxytamoxifen being the most potent metabolite^18,19^. Variability in Tam metabolism and clearance have been reported based on age, dosage, treatment duration, genetic differences, sex, and other factors, leading to different half-lives and potencies of the metabolites^19–21^. Due to these differences, there could be a concern that mice of different ages could demonstrate differences in Tam dependent cre induction that could confound studies.

To target microglia, a number of cre-ERT2 lines have been developed^22^, such as Hexb-cre^23^, P2ry12- cre^24^, Tmem119-cre^25^, Sall1-cre, and Cx3cr1-cre^26,27^. Cx3cr1-cre drivers have been the most widely used with efficient recombination and almost exclusive targeting of microglia reported^27,28^. Use of a YFP-bound Cx3cr1-cre showed up to 98% of YFP positive cells were Iba1 positive microglia with no neuronal cell labeling^27,28^. With this mouse line, there is some labeling of circulating monocytes and border associated macrophages; however, with the long life span of microglial cells, and high turnover of peripheral monocytes, labeled circulating macrophages are cleared 2-4 weeks after cre induction and the bone marrow progenitors do not demonstrate recombination^22,27–30^. For the present studies, to compare different ages of Tam induction, Cx3cr1^creERT^^2^ ^22^ mice were crossed to mice with a floxed NuTRAP allele (Nuclear Tagging and Translating Ribosome Affinity Purification)^31^ which tags nuclei with biotin/mCherry and ribosomes with eGFP^32,33^ similar to Ribotag protocols^34^. These tags can be used for INTACT (Isolation of Nuclei TAgged in specific Cell Types)^35^ isolation of nuclei or TRAP (Translating Ribosome Affinity Purification)^36^ isolation of ribosomes to obtain cell type-specific nucleic acids (DNA and RNA, respectively) from cellularly heterogeneous tissues. The fluorescent tags can be also used for imaging, cell sorting, and flow cytometry^32,37^.

This study investigates whether the effects of treating these mice with Tam at different ages (3-6 and 20 m.o.) have an impact on cre efficiency and specificity, and ability to detect age-related translatome changes in Cx3cr1-cre;NuTRAP mice^32^. We have used this experimental design of TRAP and INTACT and a floxed NuTRAP mouse model to better understand the transcriptional and epigenetic changes with age in various cell types^38–40^. The key findings from this study are that age of Tam induction had no significant effect on the efficiency or specificity of cre recombinase induction, isolation of microglial specific translatome profiles, or the ability to identify age-related microglial translatomic alterations. These results demonstrate that experimental designs utilizing early-life and late-life Tam administration are valid approaches for microglial studies and most likely generalizable to other inducible cre mouse models.

## Methods

### Mice

All animal procedures were approved by the Oklahoma Medical Research Foundation (OMRF) Institutional Animal Care and Use Committee. Breeder mice were purchased from Jackson Laboratory (Bar Harbor, ME), bred, and housed at the OMRF, under pathogen free conditions in a HEPA barrier environment. Cx3cr1^Jung-Cre/ERT2+/+^ (stock #021160)^22^ males were mated with NuTRAP^flox/flox^ females (stock # 029899)^31^ to generate the desired progeny, Cx3cr1^Jung-cre/ERT2+/wt^; NuTRAP^flox/wt^, as previously described^32^. Intraperitoneal (IP) injections of tamoxifen (Tam) solubilized by sonication in 100% sunflower seed oil (100 mg/kg body weight, 20 mg/ml stock solution, #T5648; Millipore Sigma, St. Louis, MO) were administered daily for five consecutive days^32^ at the specified ages. Tam induction at 3 m.o. and older does not lead to long-lasting changes in brain gene expression^41^ and avoids potential confounds of early post-natal Tam administration in microglial studies^42^. Based on an average weight of 20 g per mouse, each Tam injection consisted of 100 μl of 20 mg/ml stock solution. Adjustments were made for mice that significantly deviated from the average weight. Mice were euthanized by cervical dislocation, followed by rapid decapitation, in line with the AVMA Guidelines for the Euthanasia of Animals. Tam induction was performed at early (3-6 m.o.) or late age (20-21 m.o.).

### Mouse Genotyping

DNA was extracted from mouse ear punch samples by incubating samples in 50 nM NaOH at 95°C for 1 hour, after which the reaction was neutralized by adding 30 uL 1 M Tris HCl (pH: 7.4). Genotyping was performed by touchdown PCR (94°C hotstart for 2 min, 10 cycles of touchdown (94°C 20 sec, 65°C 15 sec (-0.5C per cycle decrease per cycle)), 68°C,10 sec) followed by 28 cycles of amplification (94°C 15 sec, 60°C 15 sec, 72°C 10 sec) with the listed primer sets (**Supplemental Table 1**).

### TRAP isolation

As previously described^32^, minced hippocampal tissue was Dounce homogenized (#D8938; MilliporeSigma) in 1.5 ml chilled homogenization buffer (50 mM Tris, pH 7.4; 12 mM MgCl_2_; 100 mM KCl; 1% NP-40; 1 mg/ml sodium heparin; 1 mM DTT; 100 μg/ml cycloheximide [#C4859-1ML, MilliporeSigma]; 200 units/ml RNaseOUT Recombinant Ribonuclease Inhibitor [#10777019; Thermo Fisher Scientific]; 0.5 mM Spermidine [#S2626, MilliporeSigma]; 1× complete EDTA-free Protease Inhibitor Cocktail [#11836170001; MilliporeSigma]). Homogenates were transferred to 2 ml round-bottom tubes and centrifuged at 12,000 × *g* for 10 min at 4°C. 100 μl of supernatant was saved as “Input.” The remaining supernatant was transferred to a 2-ml round-bottom tube and incubated with 5 μg/μl of anti-EGFP antibody (ab290; Abcam) at 4°C with end-over-end rotation for 1 h. Dynabeads Protein G for IP (#10003D; Thermo Fisher Scientific) were pre-washed three times in 1-ml ice-cold low-salt wash buffer (50 mM Tris, pH 7.5; 12 mM MgCl_2_; 100 mM KCl; 1% NP-40; 100 μg/ml cycloheximide; 1 mM DTT). The homogenate/antibody mixture was transferred to the 2-ml round-bottom tube containing the washed Protein-G Dynabeads and incubated at 4°C with end-over-end rotation overnight. Magnetic beads were collected (DynaMag-2 magnet) and the unbound-ribosomes and associated RNA discarded. Beads and eGFP-bound polysomes were then washed 3X with 0.5 ml of high-salt wash buffer (50 mM Tris, pH 7.5; 12 mM MgCl_2_; 300 mM KCl; 1% NP-40; 100 μg/ml cycloheximide; 2 mM DTT). Following the last wash, 350 μl of buffer RLT (QIAGEN) supplemented with 3.5 μl 2-β mercaptoethanol (#444203, MilliporeSigma) was added directly to the beads and incubated with mixing on a ThermoMixer (Eppendorf) for 10 min at room temperature. The beads were magnetically separated and the supernatant containing the target bead-bound polysomes and associated RNA was transferred to a new tube and constitutes the positive fraction for subsequent analysis. A total of 350 μl of 100% ethanol was added to the sample and loaded onto an RNeasy MinElute column (QIAGEN). RNA was isolated using RNeasy Mini kit (#74104, QIAGEN), and quantified with a Nanodrop One^c^ spectrophotometer (#ND-ONEC-W, Thermo Fisher Scientific) and quality assessed by HSRNA ScreenTape (#5067-5579, Agilent Technologies) with a 4150 Tapestation analyzer (#G2992AA, Agilent Technologies).

### Single cell suspension

Adjacent cortex tissue from the same Cx3cr1-NuTRAP mice used for hippocampal TRAP-Seq analyses were collected for flow cytometric analyses. The tissue was rinsed in ice-cold D-PBS containing calcium, magnesium, glucose, and pyruvate (#14287-072, Thermo Fisher Scientific), minced, and placed into ice-cold gentleMACS C-tubes (#130-093-237, Miltenyi Biotec), containing 1950 μl of Enzyme Mix 1. For each reaction, Enzyme Mix 1 was created by combining 50 μl of Enzyme P with 1900 μl of buffer Z, while Enzyme Mix 2 was created by combining 10 μl of Enzyme A with 20 μl of buffer Y per reaction, with all reagents included in the Adult Brain Dissociation kit (#130-107-677, Miltenyi Biotec). Transcription and translation inhibitors were included during cell preparation to prevent ex vivo activational artifacts, as previously described^37^. Actinomycin D (#A1410, MilliporeSigma) was reconstituted in DMSO at 5 mg/ml before being aliquoted and stored at −20°C protected from light. Triptolide (#T3652, MilliporeSigma) was reconstituted in DMSO to a concentration of 10 mM before being aliquoted and stored at −20°C protected from light. Anisomycin (#A9789, MilliporeSigma) was reconstituted in DMSO to a concentration of 10 mg/ml before being aliquoted and stored at 4°C protected from light. 2 μl each of actinomycin D, triptolide, and anisomycin stocks were added to the initial Enzyme Mix 1 before dissociation for a final concentration of 5 μg/ml, 10 μM, and 10 μg/ml, respectively. Each sample had 30 μl of Enzyme Mix 2 added before being mechanically dissociated for 30 min at 37°C on the gentleMACS Octo Dissociator with Heaters (#130-096-427, Miltenyi Biotec) using the 37C_ABDK_02 program. Following enzymatic and mechanical dissociation, the C-tubes were quickly spun in a chilled (4°C) Allegra-30R centrifuge (#B08708, Beckman Coulter) with an SX4400 swinging bucket rotor to collect the samples in the bottom of the tubes. Next, samples were resuspended, passed through a pre-wet 70 μm MACS SmartStrainer (#130-110-916, Miltenyi Biotec), and collected in a 50-ml conical tube (#21008-178, VWR International). The C-tubes were washed with 10 ml of ice-cold D-PBS, and the washed volume was passed through the 70 μm MACS SmartStrainer. The cells were then pelleted by centrifugation at 300 × g for 10 min at 4°C. Following centrifugation, the supernatant was aspirated and debris was removed using the Debris Removal solution (#130-109-398, Miltenyi Biotec) provided in the Adult Brain Dissociation kit (#130-107-677, Miltenyi Biotec). Briefly, cells were resuspended in 1.55 ml of ice-cold D-PBS and passed to a 5-ml round bottom tube (#22171606, FisherScientific), and 450 μl of cold Debris Removal solution was mixed into the cell suspensions. Next, 2 ml of D-PBS was gently overlaid on the cell suspension, ensuring the layers did not mix. Centrifugation at 3000 × g for 10 min at 4°C separated the suspension into three phases, of which the top two phases were aspirated. The cell pellet was gently resuspended in 5 ml of ice-cold D-PBS before centrifugation at 1000 × g for 10 min at 4°C. After aspirating the supernatant completely, the cells were resuspended in 1 ml 0.5% BSA (#130-091-376, Miltenyi Biotec) in D-PBS and filtered through a 35-μm filter (#352235, Fisher Scientific).

### RNA-Seq

Directional RNA-Seq libraries (NEBNext Ultra II Directional RNA Library, New England Biolabs, Ipswich, MA NEB#E7760) were made according to the manufacturer’s protocol as previously^37^. Poly- adenylated RNA was captured using NEBNext Poly(A) mRNA Magnetic Isolation Module (#NEBE7490). Following mRNA capture, mRNA was eluted from the oligo-dT beads and fragmented by incubating with the First Strand Synthesis Reaction buffer and Random Primer Mix (2×) from the NEBNext Ultra II Directional Library Prep Kit for Illumina (#NEBE7760; New England Biolabs) for 15 min at 94°C. First and second strand cDNA synthesis were performed sequentially, as instructed by the manufacturer’s guidelines. After purification of double-stranded cDNA with 1.8× SPRISelect Beads (#B23318, Beckman Coulter), purified cDNA was eluted in 50 μl 0.1× TE buffer and subjected to end prep. The NEBNext adaptor was diluted 1:100 in Adaptor Dilution buffer (provided) before ligating the adaptor to the cDNA. After purifying the ligation reaction with 0.9× SPRISelect Beads (#B23318, Beckman Coulter), cDNA was eluted in 15 μl of 0.1× TE (provided). Next, cDNA libraries were amplified with 16 cycles of PCR using the NEBNext Ultra II Q5 Master Mix (provided) and unique index primer mixes from NEBNext Multiplex Oligos for Illumina Library (#E6609L, New England Biolabs). Libraries were purified with 0.9× SPRISelect Beads (#B23318, Beckman Coulter) and then sized with HSD1000 ScreenTapes (#5067-5584; Agilent Technologies). Libraries had an average peak size of 316Lbp. Libraries were quantified by Qubit 1× dsDNA HS Assay kit (#Q33230, Thermo Fisher Scientific). The libraries for each sample were pooled at 4 nm concentration and sequenced using an Illumina NovaSeq 6000 system (S4 PE150). The entirety of the sequencing data are available for download in FASTQ format from NCBI Gene Expression Omnibus (GSE241574, GSE233400).

### Flow cytometry

For flow cytometric analysis, cell preparations were analyzed on a MACSQuant Analyzer 10 Flow Cytometer, as previously described^37^. Cells were stained with CD11b-APC (M1/70, #130-113-793, Miltenyi Biotec) and CD45-VioBlue (REA737, #130-110-802, Miltenyi Biotec). Following staining, cells were resuspended in 250 μl of 0.5% BSA/D-PBS. Data were analyzed using MACSQuantify v2.13.0 software.

### Immunohistochemistry

For immunohistochemistry (IHC), mouse brains were hemisected and processed for cryosectioning, as previously described^32,37^. Samples were fixed for a duration of 4h in 4% PFA, cryoprotected by overnight incubation in PBS containing 30% sucrose, and then frozen in Optimal Cutting Temperature medium (Tissue-Tek, #4583). Ten µm-thick sagittal sections were cryotome-cut and sections containing the hippocampus at the level of dentate gyrus were collected (Cryostar NX70, ThermoFisher Scientific). Tissue sections were rinsed with PBS containing 1% Triton X-100, blocked for 1h in PBS containing 10% normal donkey serum, and processed for fluorescence immunostaining. The primary antibodies included rabbit anti-mCherry (#ab167453, 1:500, Abcam, Cambridge, MA) and rat anti-CD11b (#C227, 1:200, Leinco Technologies, St. Louis, MO). Sequential imaging of brain samples was performed on an Olympus FluoView confocal laser-scanning microscope (FV1200; Olympus; Center Valley, PA) at the Dean McGee Eye Institute (DMEI) imaging core facility at the Oklahoma University Health Sciences Center (OUHSC). Microscope and FLUOVIEW FV1000 v1.2.6.0 software (Olympus) settings were identical for all samples at same magnification.

The Z-stack generated was achieved at 1.26 µm step size with a total of 6 optical slices at 40X magnification (1.2X zoom). Raw images used for figure assembly were processed using Adobe Photoshop (v24.7.0).

### Data Analysis

Following sequencing, reads were trimmed and aligned before differential expression statistics and correlation analyses in Strand NGS software package (v4.0; Strand Life Sciences). Reads were aligned against the full mm10 genome build (2014.11.26). Alignment and filtering criteria included the following: adapter trimming, fixed 2-bp trim from 5′ and 2-bp from 3′ ends, a maximum number of one novel splice allowed per read, a minimum of 90% identity with the reference sequence, a maximum 5% gap, and trimming of 3′ ends with QL<L30. Alignment was performed directionally with Read 1 aligned in reverse and Read 2 in forward orientation. All duplicate reads were then removed. Normalization was performed with the DESeq2 algorithm. Transcripts with an average read count value >20 in at least 100% of the samples in at least one group were considered expressed at a level sufficient for quantitation per tissue and those transcripts below this level were considered not detected/not expressed and excluded, as these low levels of reads are close to background and are highly variable. For statistical analysis of differential expression with aging a One-Way ANOVA with Benjamini–Hochberg multiple testing correction (BHMTC) was performed with SNK pairwise post-hoc testing. For those transcripts meeting this statistical criterion, a fold change >|1.25| cutoff was used to eliminate those genes which were statistically significant but unlikely to be biologically significant and orthogonally confirmable because of their very small magnitude of change. Visualizations of hierarchical clustering and principal component analyses (PCAs) were performed in Strand NGS (version 4.0). Gene set enrichment analysis (GSEA) was performed using GSEA v4.3.2. Heatmaps of the GSEA enrichment scores were made using Morpheus software (https://software.broadinstitute.org/morpheus).

## Results

### Microglial cell labeling specificity and efficiency are equivalent despite the age of cre induction

Single cell suspensions from aged (21-22 m.o.) cortex were collected from mice induced with Tam at either early (3-6 m.o., n=6, 4 females, 2 males) or late (20 m.o., n=7, 5 females, 2 males) age (**Figure 1A**) and stained for microglial markers CD11b and CD45. Regardless of induction age, ∼20% of all cell singlets were eGFP^+^ (**Figure 1B**). Over 95% of eGFP^+^ cells were CD11b/CD45 double positive, (**Figure 1C**) indicating that the cre recombination was equally specific for microglia at both ages of induction. One caveat of using CD45 as a microglial marker is that other resident myeloid cell types, such as macrophages, have surface expression of CD11b and CD45; however, there is the distinction between microglia and macrophages based on surface expression level of CD45^43–45^. Cells were further gated by CD11b^+^ CD45^mid^ and CD11b^+^ CD45^high^, with both induction ages having similar percentages (<10%) of CD45^high^ eGFP^+^ cells, demonstrating an equally microglial specific recombination (**Figure 1D**). No differences in the CD45^mid^ eGFP^+^ cell population by age of induction was observed. To compare efficiency of cre induction, the inverse gating strategy was used where the CD11b^+^ CD45^+^ singlets were assessed for eGFP positivity (**Figure 1E**) with the vast majority of CD11b^+^ CD45^mid^ cells being eGFP^+^, which were consistent with previously reported efficiency^46^. At greater detail, using the efficiency of eGFP labeling as a surrogate for cre recombination efficiency, of the CD11b^+^ CD45^mid^ cells, ∼90% were eGFP^+^, while in CD11b^+^ CD45^high^ cells, ∼60% were eGFP^+^ indicating that recombination efficiency is less in these macrophages (**Figure 1F**). Extending these analyses, IHC analysis of brain tissue sections in the hippocampus showed co-localization of both NuTRAP reporter proteins, eGFP and mCherry in CD11b^+^ cells (**Figure 2**), consistent with microglial identity. The pattern of recombination was undistinguishable between samples of mice subjected to early or late Tam induction. Together, these data suggest that Tam-induction of Cx3cr1-NuTRAP mice during early or late age results in similar cell specificity and efficiency of cre recombination in microglia.

**Figure 1:**
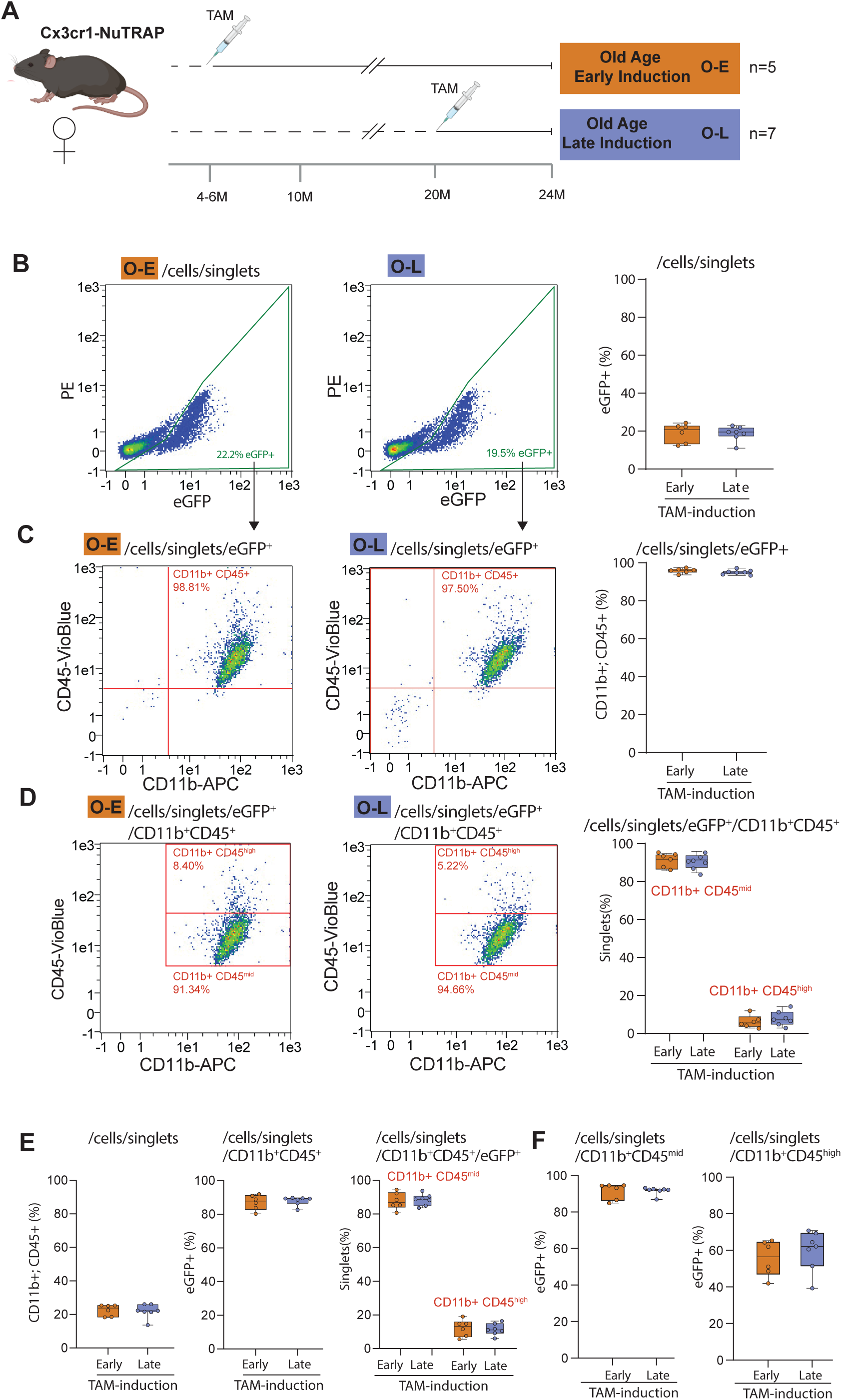
Cell labeling and efficiency are equivalent regardless of TAM induction age. A) Experimental design of early Tam induction (4-6 m.o.) and late Tam induction (20 m.o.) in Cx3cr1-NuTRAP mice followed by brain collection for both groups at 21-22 m.o. B) Single cell suspensions generated by enzymatic and mechanical dissociation of half brain cortex were gated on eGFP+ singlets, with ∼20% observed in both young and old induction mice. C) Of these eGFP+ singlets nearly all (∼98%) were double positive for CD11b and CD45. D) Further dividing this triple positive population (CD11b^+^/CD45^+^/EGFP^+^) into prototypical microglia (CD45^mid^/CD11b^+^) and other resident macrophages (CD45^high^/CD11b^+^) revealed equivalent populations. E) Using the same data, the inverse analysis was performed to determine percentage of all cells CD11b^+^/CD45^+^, of which nearly all were eGFP^+^, the vast majority of which were CD11b^+^/CD45^+^/EGFP^+^. F) ∼90% percent of CD11b+CD45^mid^ singlets were eGFP+, indicative of high efficiency of cre recombination for microglia in our model with both ages of Tam induction. Similar efficiency of recombination was observed at both ages of induction when analyzing %eGFP in CD45^high^/CD11b^+^ cells (putative macrophages)

**Figure 2.**
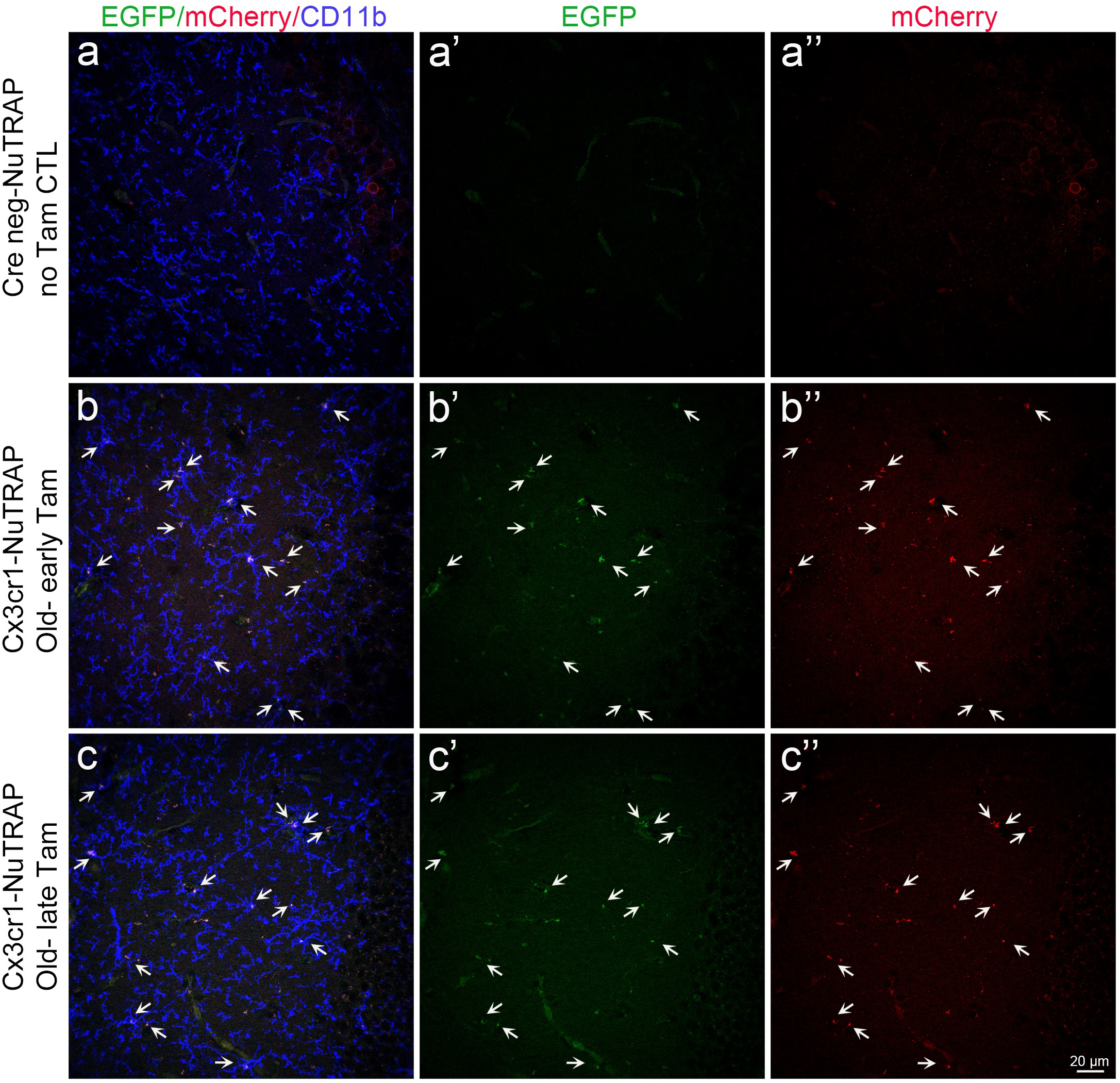
DNA Recombination in the brain of old Cx3cr1-NuTRAP mice subjected to tamoxifen treatment during early and late life. Cx3cr1-cre^+^ NuTRAP^+^(Cx3cr1-NuTRAP) mice were treated with Tam at 4-6 m.o. (early induction) or 20 m.o. (late induction). At 21-22 m.o., brains were harvested and processed for immunohistochemical analysis of microglial recombination. Frozen tissue sections were immunostained with antibodies against mCherry and CD11b. Representative confocal fluorescent microscopy images of sagittal brain sections show that unlike cre negative counterparts (controls) (a-a’’), Cx3cr1-NuTRAP brains displayed colocalization of eGFP (green signal), mCherry (red signal), and CD11b (blue signal) following early (b-b’’) and late (c-c’’) Tam administration regimes. The area imaged for each sample encompasses the dentate gyrus of the hippocampus. The scale bar depicts 20 µm.

### Age of cre induction does not alter TRAP isolation of microglial-specific RNA efficiency

Once establishing specificity and efficiency of recombination of the NuTRAP allele at both ages of Tam induction, we set out to determine if the downstream TRAP isolation of RNA demonstrated similar profiles. The NuTRAP allele labels ribosomes with eGFP, which can then be isolated to collect cell type-specific RNA for further analysis^32^ (**Figure 3A**). To determine the consistency of microglial specific RNA enrichment at different timepoints of Tam administration, mice were treated at an early age (3 m.o) and collected at young age (4 m.o., n=5, 5 female, Young Age – Early Induction) or were tram treated early (at 3-6 m.o, n=7, 7 female, Old Age – Early Induction) or late (at 20 m.o., n=5, 5 female, Old Age – Late Induction) and hippocampi collected at old age (21-22 m.o.). Samples were processed by TRAP and input and positive fraction RNAs collected for RNA seq analysis (**Figure 3A**). Principal component analysis (PCA) of all expressed genes showed separation by fraction (positive or input) in the first component (67.6% explained variance), but there was no further separation by age of induction or age of collection (**Figure 3B**). To determine if the timing of Tam induction resulted in similar enrichment of hippocampal microglial translatomes, we compared enrichment (positive fraction/input fraction) of marker genes of various brain cell types^47^ following early and late Tam induction. We further compared this data to previously obtained RNA sequencing data from sorted microglial cells as a positive control. TRAP isolated RNA demonstrated equivalent enrichment for microglial marker genes, regardless of induction or collection age, and almost identical enrichment levels as compared to sorted microglial cells (**Figure 3C**). Additionally, there was depletion of marker genes for astrocytes, neurons, and oligodendrocytes in the positive fraction as compared to the input (**Figure 3C & E**). Statistical comparisons are detailed in Supplemental Table 2. The fold enrichment/depletion for all of these cell type marker genes demonstrated correlations over 0.97 for each age of Tam induction and age of collection (**Figure 3D**). Overall, these findings demonstrate that microglial RNA enrichment is not affected by the age of Tam induction or collection.

**Figure 3:**
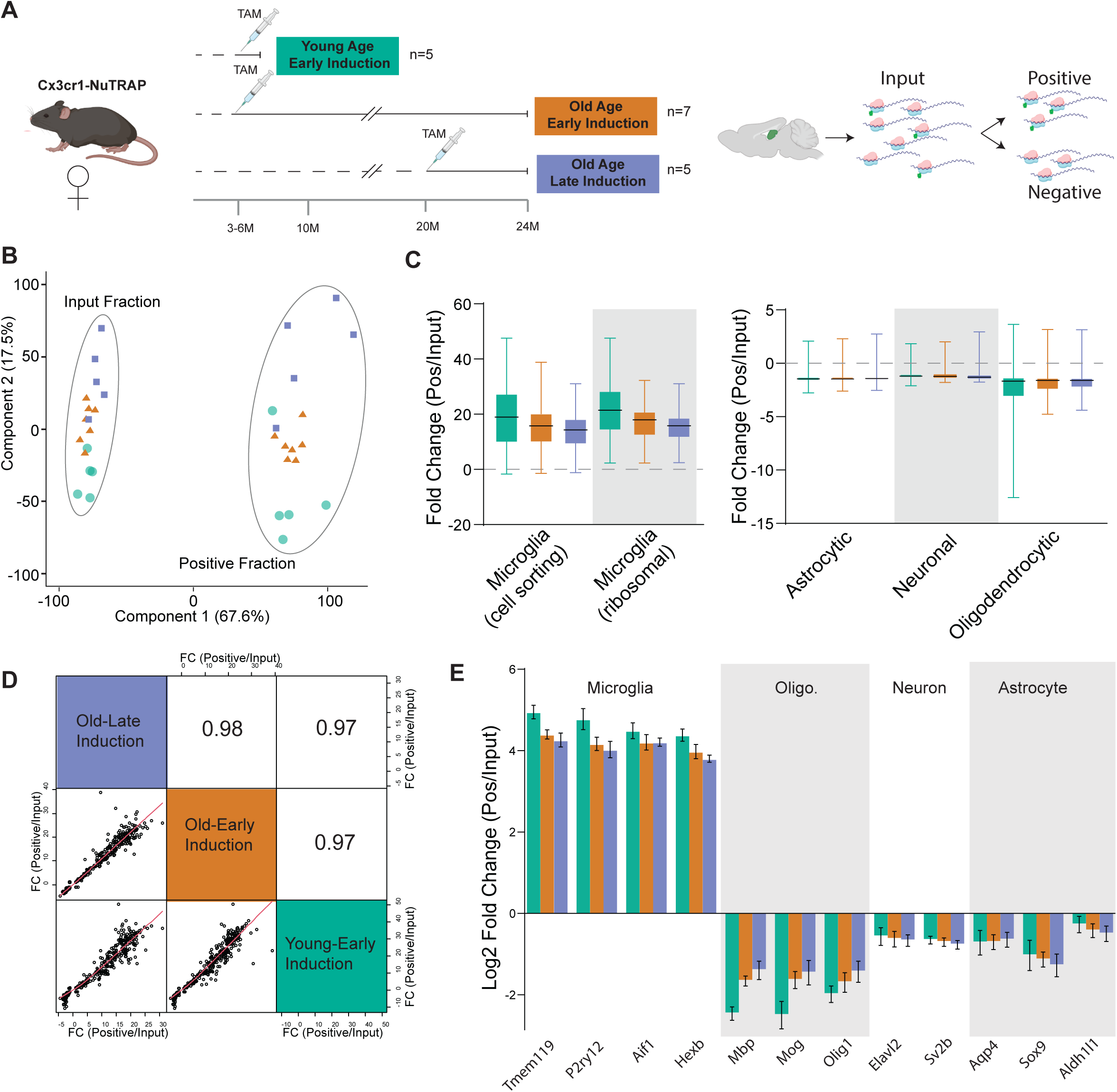
Translatomes from young and old inductions are equivalently enriched for microglial markers. A) Cx3cr1-NuTRAP mice were induced at 3-6 m.o. and collected at 4 m.o. or 21-22 m.o. or induced at 20 m.o. and collected at 21-22m.o. TRAP isolation was performed on hippocampal dissections and RNASeq profiles generated from Input and Positive fractions. B) Translatome profiles separated in the first component between the Input and Positive fractions in the first component and to a degree between young and old mice in the 2^nd^ component. C) Assessment of CNS cell type markers derived from cell sorting and ribosomal profiling studies revealed an equivalent enrichment of microglial markers and depletion of markers for other cell types. D) Correlation analysis of the union of marker lists demonstrated a high congruency of enrichment (fold change Positive fraction/Input fraction) across ages and ages of TAM induction. E) Example marker gene enrichments/depletion for microglia, neuron, astrocyte, and oligodendrocyte markers.

### Age of Tam induction does not affect the trajectory of age-related changes in the brain

Once NuTRAP allele induction was determined to be equivalent in both ages of Tam induction and collection, we asked whether timing of Tam induction can affect biological endpoints of importance. We and others have identified significant transcriptional changes in hippocampal microglia with aging^40,48,49^. We compared gene expression data from the TRAP positive fractions of old mice following early Tam induction (n=7, 7 females) and old mice following late Tam induction (n=5, 5 females) to young mice following early Tam induction (n=5, 5 females) to analyze differential gene expression (One way ANOVA, Benjamini Hochberg Multiple Testing Correction q<0.1, Fold Change >|1.25|) with aging. As visualized by hierarchical clustering, the translatomic differences with age of the two old groups are similar to each other but differ from those of the young group (**Figure 4A**). This supports that the differential gene profiles with aging of the two old groups are not being significantly affected by age of induction. Differential expression with aging in the old early and late induction as compared to young were strongly correlated, with nearly all differentially expressed transcripts demonstrating the same directionality and similar magnitude of change. (**Figure 4B**). Since a few genes have an inverse fold change trend on the plot, we created boxplots of these genes (**Supplemental Figure 1**). The boxplots from these genes demonstrate minor changes between the old early induction and old late induction that may represent random variability. These findings confirm that the cre induction is not only similar with various ages of induction, but that induction in late age is not inducing confounds and in fact similar populations of microglia are being examined. This supports that different induction ages do not induce confounding variables when using age as an endpoint.

**Figure 4:**
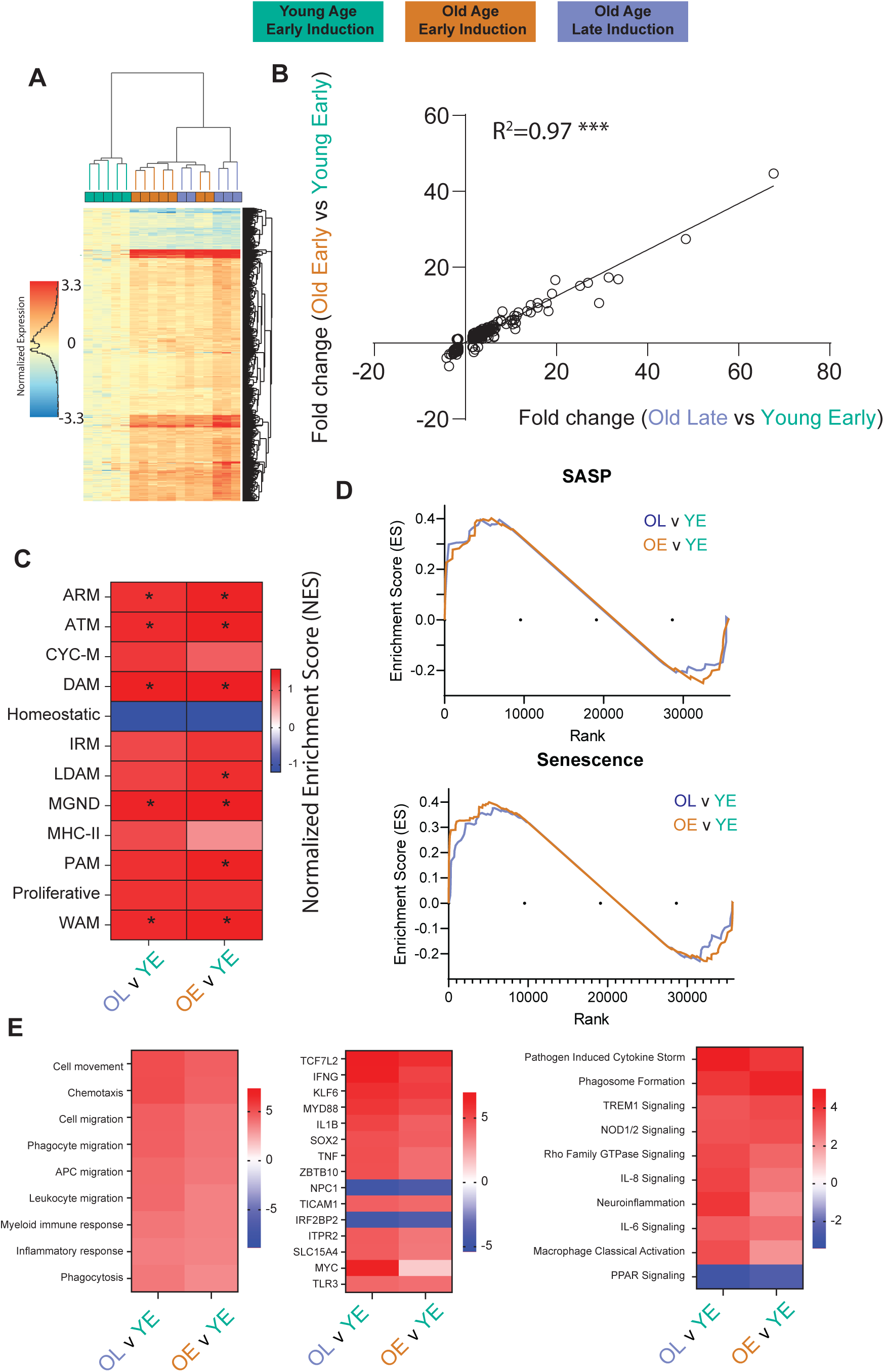
Differentially expressed genes with age for different ages of induction. A) Heat map of differentially expressed genes comparing both old early induction and old late induction to young early induction mice. The heatmap demonstrates similarity in differential expression in both old groups, allowing for equivalent detection of age-related changes regardless of induction age. B) Comparing the age-related fold changes of differentially expressed genes from the different induction ages correlate highly between both ages of induction. C) Heatmap of the GSEA enrichment scores comparing enrichment of microglial phenotypic markers (Supplementary Table 3, gene lists) by tamoxifen induction time (OL v YE) and age (OE v YE) in positive fractions. D) GSEA enrichment plots comparing enrichment of senescence associated secretory phenotype (SASP) and senescence markers (Supplementary Table 3). E) Relative Z-score from IPA for disease pathways, upstream regulators, and canonical pathways for age related changes with various induction ages showing no difference between the old mouse groups.

Using the same data, we investigated the relative enrichment of different microglial subtypes with the different ages of induction. Microglia exhibit various phenotypic states^50–53^ in response to sterile inflammation with aging. Several subtypes of microglia have been defined, including: 1) disease-associated (DAM)^54^, 2) interferon-response (IRM)^55^, 3) lipid-droplet (LDAM)^56^, 4) white-matter-associated (WAM)^57^, 5) activated-response (ARM)^55^, 6) neurodegenerative (MGnD)^58^, and 7) proliferative-region-associated (PAM)^59^ microglia, among others. Recent comparisons have suggested that, perhaps, there are four fundamental patterns underlying these many microglial phenotypes (DAM, IFN, MHC-II, proliferating, etc.)^60^. One pattern of note is Disease-Associated Microglia (DAM), first reported^61^ and recently reviewed^62^, which are proposed to develop from homeostatic microglia through a progressive change that is initially TREM2-independent (DAM1) and then TREM2-dependent (DAM2). Investigating these various phenotypic states with age is essential to understanding the involvement of microglial inflammation with neurodegeneration, and being able to compare various induction ages of select mouse models with these endpoints is advantageous. As such, we used Gene Set Enrichment Analysis (GSEA) to compare microglial phenotypic states of our old mice at different induction ages to our young mice (**Figure 4C**) using gene sets developed from the literature (**Supplemental Table 3**). Comparing old mice to young mice, with both ages of induction, we see an enrichment for reactivated microglial subtypes, such as DAMs, ARMs, MgNDs, and WAMs, while also seeing a depletion of homeostatic microglia. This is consistent with previous findings by our labs and others^40,63,64^, and there is no differential effect due to age of induction.

In addition to microglial phenotypic changes, microglial senescence is emerging as a contributing factor to inflammaging^65,66^. Evidence demonstrates that senescent microglia contribute to cognitive decline and white matter degradation^67^, and this pattern is observed in multiple different microglial subtypes and disease progressions^40,63,68^. We used similar GSEA approaches with senescent cell marker lists (**Supplemental Table 3**) to detect enrichment of senescent cells in our aged mice. Both senescent cell type and senescence-associated secretory phenotype (SASP) were enriched in our old mice compared to young, with equivalent enrichment at both induction ages (**Figure 4D**). This coincides with current research and further supports the effectiveness of using different induction ages of cre models to investigate various aspects of aging in mice. We further analyzed this data through Ingenuity Pathway Analysis, and there were observed changes in disease pathways, canonical pathways, and upstream regulators associated with aging that were similar regardless of induction age (**Figure 4E**).

### Specific microglial responses with age are unaffected by age of Tam induction

For a more focused analysis of the specific changes seen with aging using the different induction ages of the cre-NuTRAP model, we identified specific cell markers for various conditions or phenotypic states of interest. While enrichment of DAMs was already determined by GSEA, focusing on the most prominent (**Figure 5A**) DAM markers, there was consistent increase in old samples, regardless of induction age. Examining homeostatic markers, which are known to reduce with age, both ages of induction demonstrated a decrease of homeostatic markers, consistent with expected findings (**Figure 5B**). Senescent cell markers demonstrated the same pattern, with no effect noted by different induction ages (**Figure 5C**). We further chose to differentiate between the p16^Ink4a^ and p19^Arf^ isoforms of Cdkn2a. p16^Ink4a^ is more associated with senescence and demonstrated a significant increase, consistent between induction ages while p19^Arf^ had smaller increases with age (**Figure 5D**). In our recent work, we have identified MHC-I components as upregulated with age in microglia from humans, mice, and rats^48^. To assess whether this result is recapitulated with the different induction ages, we analyzed expression of genes for MHC-I and their receptors with early and late Tam induction. We observed increased expression of MHC-I genes B2m, H2-D1, H2-K1, and H2-M3, as well as MHC-I receptors, Lilrb3, Lilrb4, and Pilra (**Figure 5E**). These increases were seen in both induction ages to a similar degree, demonstrating the effectiveness of different induction ages that continues to correlate with previous aging data. In a smaller set of male mice (Young n=4, old-early n=5, old-late n=2) we performed a similar analysis and found no/limited differences with age of induction (**Supplemental Figure 2**).

**Figure 5:**
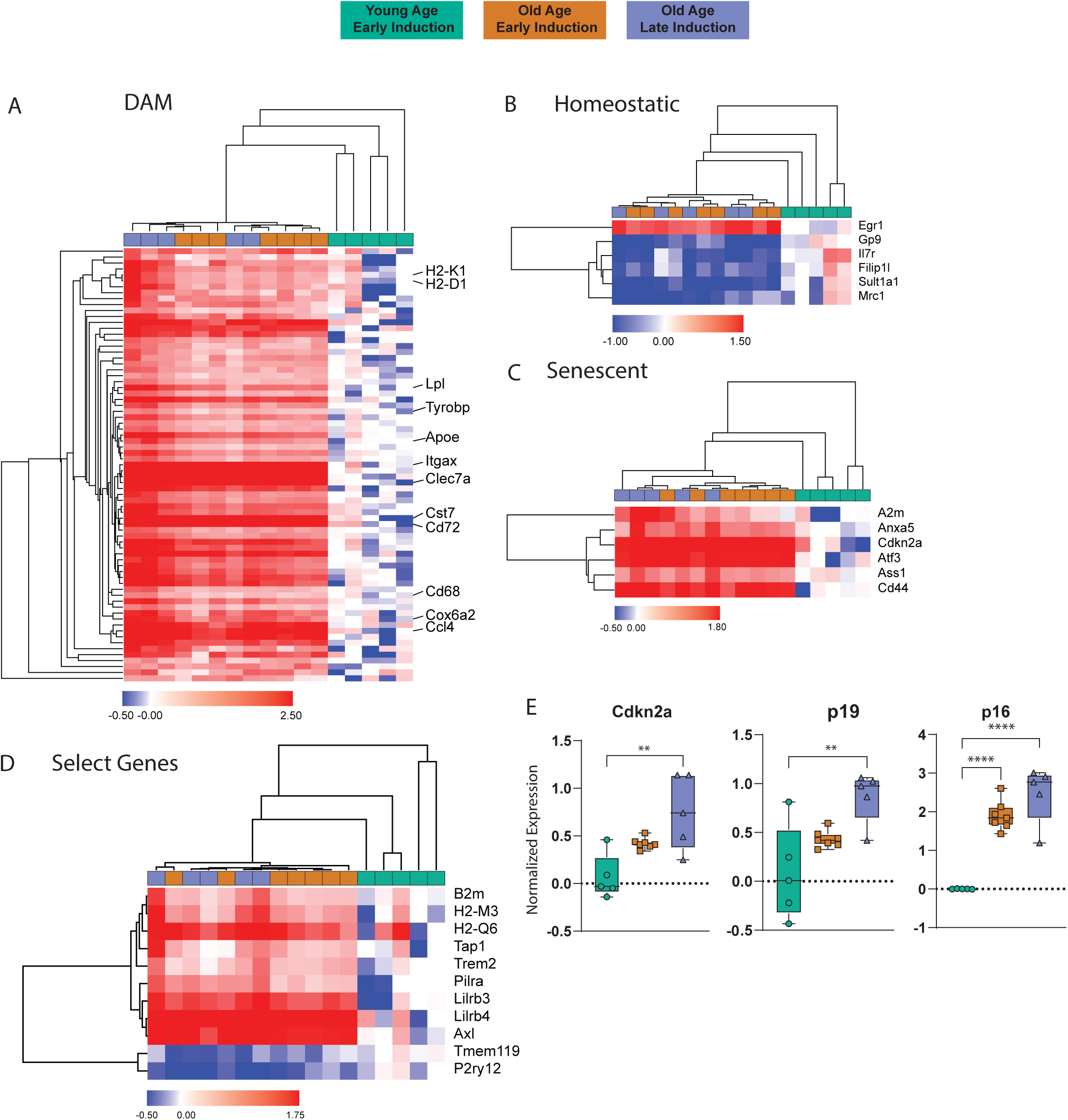
Enrichment of microglial RNA isolated from various ages of induction. A) Heatmap created from markers for DAMs demonstrating similar enrichment of individual genes for DAMs regardless of age of induction. B) Heatmap for homeostatic microglia showing consistent downregulation in both old age groups. C) Heatmap for senescence related genes were equivalently upregulated regardless of the age of Tam induction. D) Boxplots showing the change with age in both groups of Cdkn2a and both of its isoforms, p19^Arf^ and p16^Ink4a^. E) Select genes of interest based on other microglial aging studies showing similar patterns of differential expression regardless of age of induction.

## Discussion

The focus of this study was to evaluate and characterize the effectiveness of different ages of Tam induction in a Cre/ERT2 mouse model. A microglial NuTRAP mouse model was used to compare induction efficiency and specificity, and to capture age-related changes in the microglial translatome. Tam inducible models have emerged as a critical tool for temporally controlling transgene expression to elucidate gene function, as well as testing the therapeutic potential of different gene targets. In aging research studies, inducing cre at older ages is potentially highly useful for modulating late-life gene expression without disturbing developmental or adult gene expression. Studies have identified some difference in Tam metabolism with age, sex, and other factors^19^, and a few studies have described limitations and acute negative effects of Tam induction in embryonic and early postnatal models^23,69^. However, to our knowledge there are no studies that have empirically tested for equivalent specificity, efficiency, or consistency of detecting age-related biological changes after inducing cre recombination at older ages. We used a Cx3cr1-creERT2 driver crossed with a NuTRAP allele as a testing platform that labels microglia and allows for ribosomal isolation to analyze cell type-specific translating RNA^32^. This strategy allows for both identification of labeled cells and assessment of molecular phenotypes. We examined if the cre induction of our NuTRAP allele maintained the same efficiency and specificity at either early (3-6 m.o.) or late (20 m.o.). Using ribosomal pulldown and translatome sequencing, we then compared expression patterns and the ability to detect age-related changes.

Flow cytometry analysis of eGFP+ cells demonstrated high and equivalent levels of specificity and efficiency at both induction ages. Of note, while efficiency of induction was >90% for Cd11b+ CD45^mid^, presumably microglia, it was lower (∼60%) for Cd11b+ CD45^high^ cells which are likely other macrophage cell types. The cell labeling data provides a high degree of confidence that different ages of induction will not cause confounding effects due to differential cre efficiency or specificity.

Molecular analysis of TRAP isolated RNAs allows for an in-depth analysis of cell labeling and for comparison of age-related gene expression changes. Assessment of cell marker lists developed by the field^47^ showed equivalent enrichments of microglial markers in young and old mice, and from different induction ages. Additionally, depletion of markers for other cell types was the same between groups. This provides further evidence of similar specificity of microglial labeling across all groups. In analysis of age-related changes, highly consistent alterations in gene expression were observed in old mice as compared to young mice, regardless of the induction age of the old mice. Prototypical signatures of microglial senescence and Disease Associated Microglia (DAM) were evident in old mice. Further, we did see differences between the young and old groups in homeostatic microglial markers (e.g., P2ry12 and Tmem119) which corresponds with other studies that demonstrate a reduction in expression of homeostatic markers with age in microglia^70^. The fold changes for age-related changes for the different induction ages were nearly identical, with high correlation. The late induction group had slightly higher fold changes which may be attributable to a batch effect, as this was mainly driven by three of the samples this group collected at the same time. Previously, we have examined the potential for Tam to cause sexually divergent, long-lasting changes in gene expression^41^ but found none. We examined males and females in this study. The main findings were in female mice with the analysis of a smaller set of male samples demonstrating the equivalent molecular patterns.

With the concept of antagonistic pleiotropy in the field of Geroscience^71^, and in neurodegeneration research where prodromal versus disease progression studies are needed, the use of Tam inducible cres in older mice has many applications. For example, C1q is necessary for microglia for development and neuronal pruning, but it can become harmful later in life. Thus C1q-cre knockouts would benefit from being able to induce the knockout at various ages to better establish the different involvement of C1q at various points in neurodegeneration^72^. Additionally, with the emergence of microglial phenotype analysis, subtypes populations such as DAMs, WAMS, LDAMs, etc. increase with age and may have a protective effect that becomes deleterious^63^. Studies involving these genes and subtype analysis will benefit from changing the in vivo genome at different ages to visualize the effects that can only be elucidated in an aged environment.

While these findings may be generalizable across cre/ERT2 systems, it is optimal to test the consistency of specificity, efficiency, and molecular phenotypes that results from cre induction at different ages with each cre and floxed gene combination. Mouse line specific factors (e.g., cell type, size of flox construct) could alter these results. However, these results do demonstrate that Tam inducible cre induction in older mice is a practical approach for aging studies to understand the gene function at different ages.

---

**Supplemental Table 1:** Genotyping primers for ear punches for genotype testing of mice.

**Supplemental Table 2:** Statistical significance for differences between groups on individual marker genes for cell types, relating to Figure 3.

**Supplemental Table 3:** Cell marker lists used for GSEA analysis.

**Supplemental Figure 1:** Boxplots for differentially expressed genes in aging that had an apparent inverse relationship. This figure demonstrates that the apparent inverse relationship is likely due to one of the groups having incredibly low statistical significance.

**Supplemental Figure 2:** Brief analysis performed on male mice same as female mice. This figure is included in the supplement as one of the male groups had an n of 2, which is too underpowered for a full analysis. A) GSEA for cell types showing enrichment in all groups for microglia and depletion of all other cell types. B) Enrichment for specific marker genes further showing microglial enrichment and other cell depletion. C) Age related differential genes comparing fold changes between induction ages showing high correlation. D) GSEA heat map showing similar enrichment of microglial phenotypes. E) Specific genes selected for analysis showing similar age-related changes regardless of induction age.

## Acknowledgments

The authors would like to acknowledge David Bowman for assistance with figure preparation.

