## Supplementary figures and images for "Consistent specificity and efficiency of tamoxifen-mediated cre induction across ages"

### Supplemental Figure 1

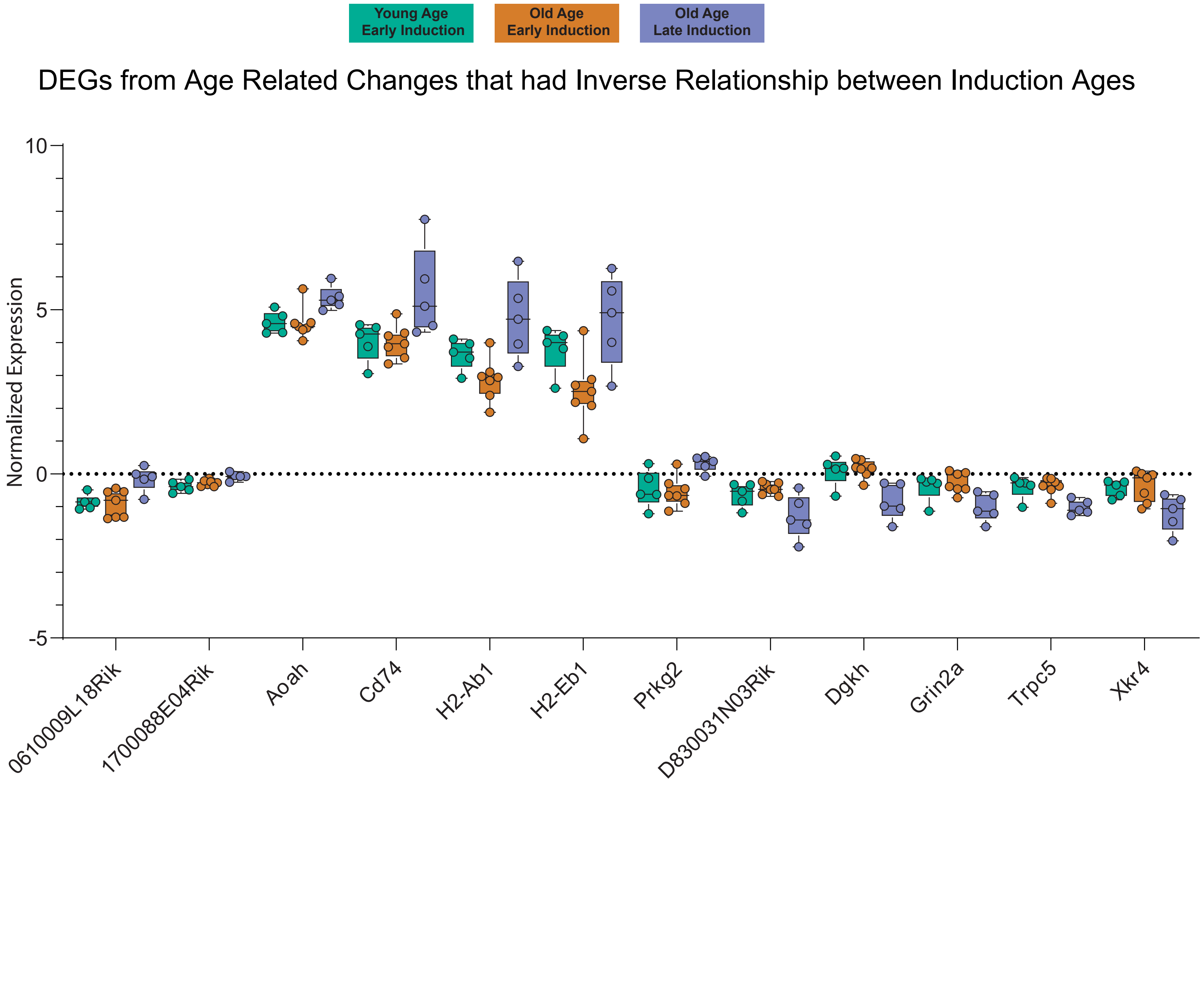

### Supplemental Figure 2

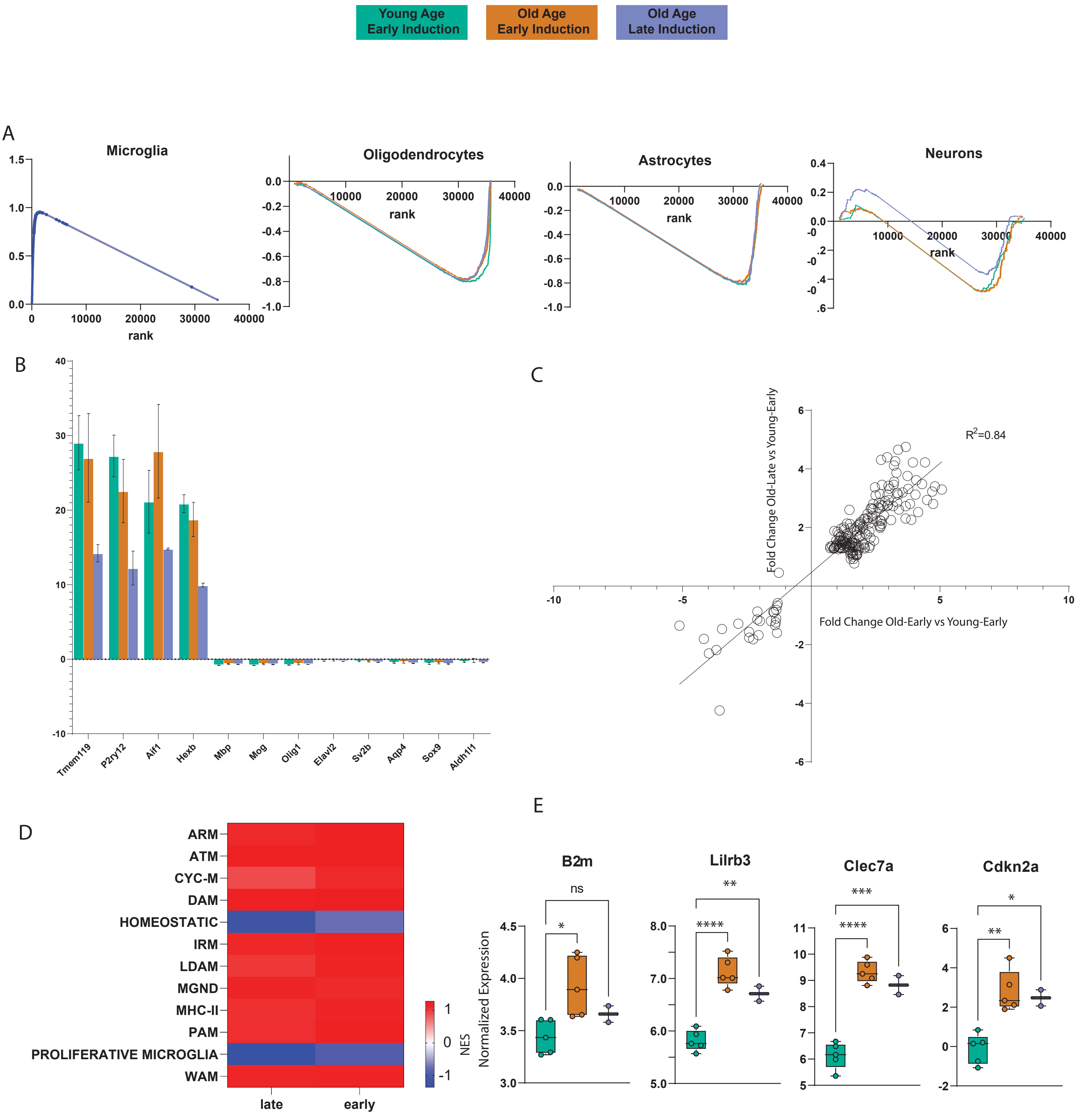
